# Step-by-step protocol for making a knock-in *Xenopus laevis* to visualize endogenous gene expression

**DOI:** 10.1101/2025.03.05.640691

**Authors:** Norie Kagawa, Yoshihiko Umesono, Ken-ichi T Suzuki, Makoto Mochii

## Abstract

We established a novel knock-in technique, New and Easy *Xenopus* Targeted integration (*NEXTi*), to recapitulate endogenous gene expression by reporter expression. *NEXTi* is a CRISPR-Cas9-based method to integrate a donor DNA containing a reporter gene (*egfp*) into target 5’ untranslated region (UTR) of *Xenopus laevis* genome. It enables us to track eGFP expression under regulation of endogenous promoter/enhancer activities. We obtained about 2% to 13% of knock-in vector-injected embryos showing eGFP signal in a tissue-specific manner, targeting *krt.12.2.L*, *myod1.S*, *sox2.L* and *bcan.S* loci, as previously reported. In addition, F1 embryos which show stable eGFP signals were obtained by outcrossing the matured injected frogs with wild-type animals. Integrations of donor DNAs into target 5’ UTRs were confirmed by PCR amplification and sequencing. Here, we describe the step-by-step protocol for preparation of donor DNA and single guide RNA, microinjection and genotyping of F1 animals for the *NEXTi* procedure.

**Highlight:**

- Intended organism: *Xenopus laevis*
- Purpose of the protocol: Efficient knock-in for visualizing endogenous target gene expression using CRISPR-Cas9 system
- Essential equipment and materials: Microinjector, fluorescence microscopy, Cas9 protein, sgRNA, donor DNA
- Features: Expression of reporter genes depends on endogenous enhancer/promoter activities.

## 2. Background and Features

Targeted integration or knock-in of a donor construct into a desired locus is a powerful tool which enables visualization of endogenous gene expression or manipulation of gene function in both model organisms and non-model organisms including amphibians. We previously established a simple transgenic method called New and Easy *Xenopus* Transgenesis (*NEXTrans*) in *Xenopus laevis* (Shibata et al., 2022; Shibata et al., 2023), by which the donor plasmid was efficiently integrated into the *tgfbr2l* loci, the proposed safe harbor sites, through the non-homologous end joining (NHEJ) repair induced by Cas9 ribonucleoprotein (RNP). Using a similar strategy to *NEXTrans* we recently succeeded in the targeted integrations of a reporter gene, an enhanced green fluorescent protein (eGFP) gene, into 5’ UTRs of various target genes to enable visualization of endogenous genes’ expression and termed the method New and Easy *Xenopus* Targeted integration (*NEXTi*) (Mochii et al., 2024). In the *NEXTi* procedure, fertilized eggs were injected with Cas9, a single guide RNA (sgRNA) for the 5’ UTR of the target gene and a donor plasmid containing *egfp* flanked by the target genomic sequence (Fig. 1A). The donor *egfp* was integrated into the target locus through essentially the NHEJ process and was expressed under control of the endogenous promoter/enhancer activity (Fig. 1B). This versatile method can be applied for targeted integrations into various genome regions and will contribute much to developmental and regeneration studies in *Xenopus*.

**Fig. 1.**
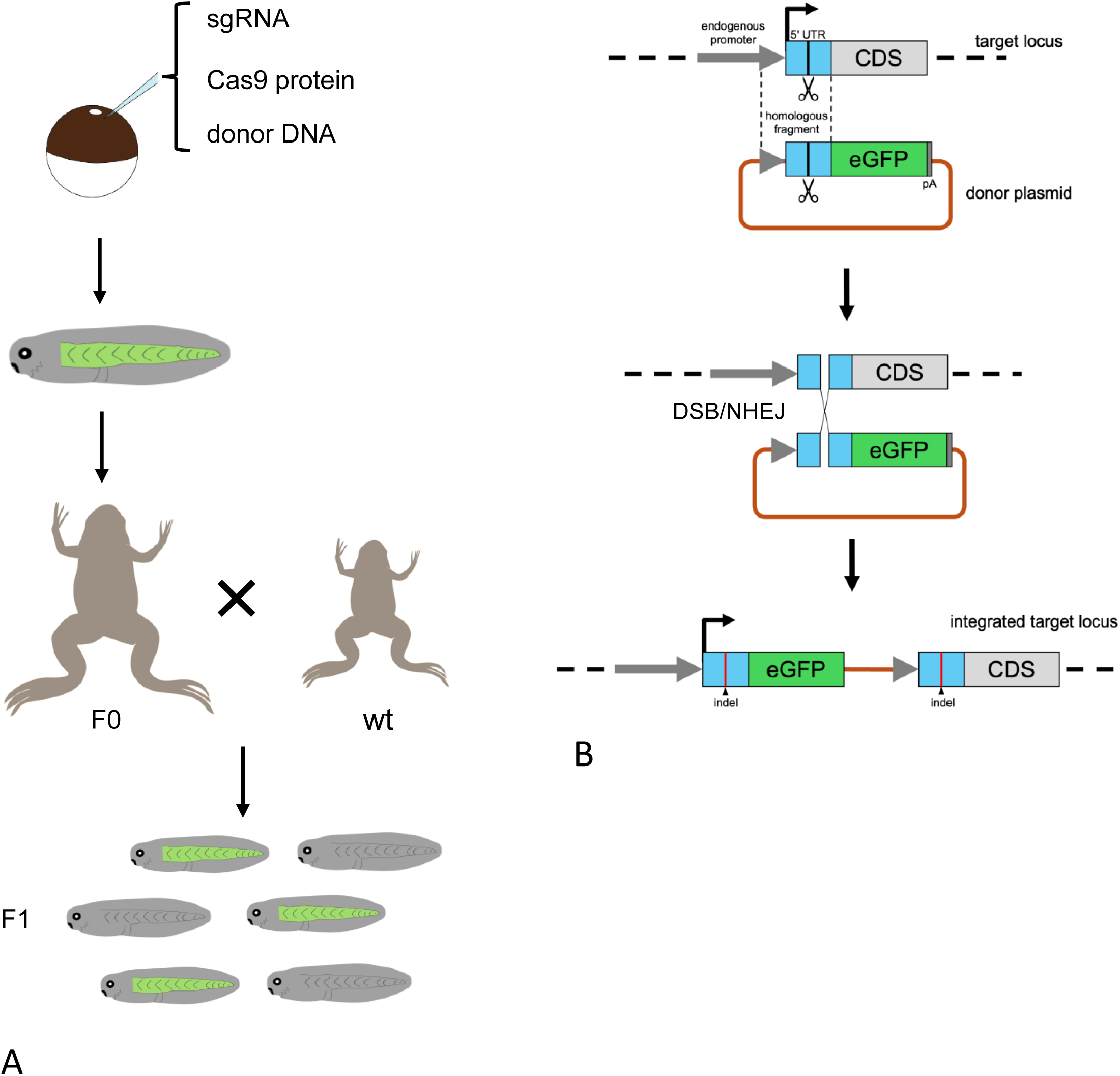
Overview of *NEXTi*. (A) Summary of the *NEXTi* procedure to make a knock-in founder and F1 offsprings. The *NEXTi* donor plasmid carrying the *egfp* is injected into fertilized *Xenopus laevis* eggs with Cas9 protein and a single guide RNA. Selected founder (F0) expressing tissue-specific eGFP is outcrossed with wild-type animal (wt) to obtain F1 animals. (B) Schematic representation of targeted integration by *NEXTi.* The Cas9 RNP binds and cleaves both the target sequences in the donor plasmid and the 5’UTR of the target locus. The plasmid integrates into the target sites in the zygotic genome via DSB/NHEJ. Insertion and/or deletion (indel) are found in the upstream and downstream junctions.

### Advantages

*NEXTi* enables faithful recapitulation of the endogenous gene expression and can be applied for any target gene in *X. laevis*. As sgRNA target sequence is located within the 5ʹ UTR of the target locus in this method, some insertions and/or deletions (indel) of nucleotide at the junction between the target sequence and injected donor usually do not disturb expression of the reporter gene. Therefore, *NEXTi* is superior to knock-in methods targeting coding regions in which indels often causes serious effects on the reporter expression.

### Limitations

As the donor integration is mediated essentially through the NHEJ process, orientation of the integration is not controlled and probably random in *NEXTi*. The reporter gene integrated in reverse orientation will be expressed incorrectly. The donor DNA should not contain the core enhancer of the target gene locus. Such an enhancer would induce reporter expression mimicking endogenous expression even in off-target insertion. Genes with a short 5ʹ UTR could not be targeted by *NEXTi*, since an sgRNA could not be designed within the short sequence. Integration of the donor within the 5’UTR will interfere normal expression of the targeted allele and may disturb normal development. As the efficiency of the targeted integration varies among different contexts, it may be necessary to try multiple sgRNAs for a single locus in order to achieve successful integration.

### Applications

As mosaic expression of the reporter gene is frequently observed in the founders, we obtained F1 offspring with stable reporter expression in multiple lines (Mochii et al., 2024; this work). The *NEXTi* strategy can be applied for targeted integrations into various genomic regions other than 5’UTR. For example, integration of a modified exon into an intron will result in a replacement of the target coding sequence with a modified one.

## 3. Materials and Equipment

### 3.1 Solution

10 × Marc’s Modified Ringers (MMR) stock solution

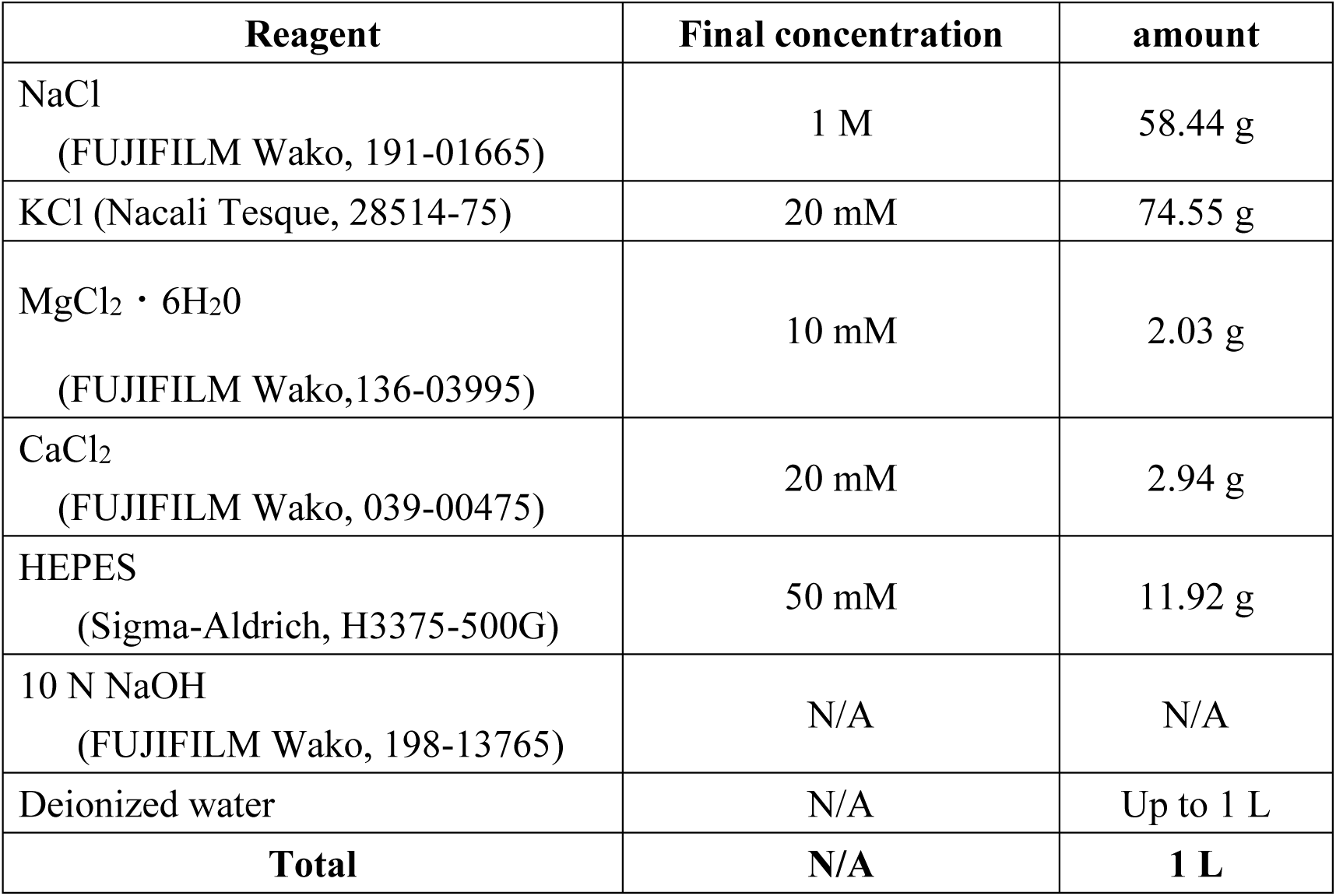

| Reagent | Final concentration | amount |
| --- | --- | --- |
| NaCl<br>(FUJIFILM Wako, 191-01665) | 1 M | 58.44 g |
| KCl (Nacali Tesque, 28514-75) | 20 mM | 74.55 g |
| MgCl<sub>2</sub> · 6H<sub>2</sub>O<br>(FUJIFILM Wako, 136-03995) | 10 mM | 2.03 g |
| CaCl<sub>2</sub><br>(FUJIFILM Wako, 039-00475) | 20 mM | 2.94 g |
| HEPES<br>(Sigma-Aldrich, H3375-500G) | 50 mM | 11.92 g |
| 10 N NaOH<br>(FUJIFILM Wako, 198-13765) | N/A | N/A |
| Deionized water | N/A | Up to 1 L |
| **Total** | **N/A** | **1 L** |

Adjust the solution to pH 7.5 with 10 N NaOH.

0.2% Ethyl 3-aminobenzoate methanesulfonate (MS222)

Adjust the solution to pH 7.5 with 10 N NaOH. 0.2% Ethyl 3-aminobenzoate methanesulfonate (MS222)
| Reagent | Final concentration | Amount |
| --- | --- | --- |
| MS222<br>(Sigma-Aldrich, A5040-100G) | 0.2% (w/v) | 2 g |
| NaHCO<sub>3</sub><br>(FUJIFILM Wako, 191-01305) | 16.7 mM | 1.4 g |
| Deionized water | N/A | Up to 1 L |
| **Total** | **N/A** | **1 L** |

De-jellying solution

| Reagent | Final concentration | Amount |
| --- | --- | --- |
| L-cysteine<br>(FUJIFILM Wako, 037-05275) | 2% (w/v) | 4 g |
| 10 N NaOH<br>(FUJIFILM Wako, 198-13765) | N/A | N/A |
| Deionized water | N/A | Up to 200 mL |
| **Total** | **N/A** | **200 mL** |

Adjust the solution to pH 8.0 with 10N NaOH.

10x Cas9 buffer

Adjust the solution to pH 8.0 with 10N NaOH. 10x Cas9 buffer
| Reagent | Final concentration | Amount |
| --- | --- | --- |
| 2 M KCl<br>(Invitrogen, AM9640G) | 1.5 M | 15 mL |
| 1 M HEPES (pH 7.5)<br>(Nacalai Tesque, 15639-84) | 0.2 M | 4 mL |
| Nuclease-free water | N/A | 1 mL |
| (Nacalai Tesque, 06442-95) |  |  |
| **Total** | **N/A** | **20 mL** |

3% Ficoll-0.5 x MMR (-0.1 x MMR)

| **Reagent** | **Final concentration** | **Amount** |
| --- | --- | --- |
| 10 × MMR | 0.5 × MMR<br>(0.1 × MMR) | 25 mL<br>(5 mL) |
| Ficoll PM400 (Sigma-Aldrich,<br>F4375)<br>(GE Healthcare, 17-0300-50) | 3% (w/v) | 15 g |
| Kanamycin Sulfate<br>(FUJIFILM Wako, 113-00343) | 20 ng/mL | 10 mg |
| Deionized water | N/A | up to 500 mL |
| **Total** | **N/A** | **500 mL** |

0.3% methyl-cellurose-0.5 x MMR (-0.1 x MMR)

0.3% methyl-cellulose-0.5 x MMR (-0.1 x MMR)
| **Reagent** | **Final concentration** | **Amount** |
| --- | --- | --- |
| 10 × MMR | 0.5 × MMR<br>(0.1 × MMR) | 25 mL<br>(5 mL) |
| Methyl Cellulose 400<br>(FUJIFILM Wako, 132-05055) | 0.3% (w/v) | 1.5 g |
| Kanamycin Sulfate<br>(FUJIFILM Wako, 113-00343) | 20 ng/mL | 10 mg |
| Deionized water | N/A | up to 500 mL |
| **Total** | **N/A** | **500** |

### 3.2 Setup

#### 3.2.1 Microneedle

Microneedles are made by pulling glass capillaries (Drummond Scientific, 3-000-204R) with a puller (e.g. Narishige, PC-100). The tip of the needle is sharpened and adjusted to an outer diameter of 20–25µm (See Tip1).

#### 3.2.2 Injection plate

To make an injection plate a 60 mm petri dish is covered with a modeling clay, Plastalina. Spread the clay to make it flat and make grooves in it (See Tip2).

### 3.3 Other reagents and equipment

PrimeSTAR MAX (Takara Bio, R010A)

PureYield Plasmid System (Promega, A2490)

Ex Taq Hot Start Version (Takara, RR006)

Wizard SV Gel and PCR clean-up system (Promega, A9282)

T7 RiboMax express large-scale RNA production system (Promega, P1320)

Alt-R S.p. Cas9 Nuclease 3NLS (Integrated DNA Technologies, 1081058)

Tris-HCl (Sigma-Aldrich, T5941)

EDTA (Sigma-Aldrich, E-6511)

Sodium lauryl sulfate (SDS) (Nacali Tesque, 31607-65)

Proteinase K (FUJIFILM Wako, 160-14001)

RNase A (Sigma-Aldrich, from bovine pancreas)

Serotropin (1,000 U/glass ampoule) (ASKA Animal Health)

Human Chorionic Gonadotropin (HCG) (5,000 U/glass ampoule) (ASKA Animal Health)

Fluorescent stereomicroscope (Leica, MZ165FC)

Microinjector (Drummond Scientific, 3-000-204)

Micromanipulator (Drummond Scientific, 3-000-204R)

PCR thermal cycler (ASTEC, G02)

NanoDrop One (Thermo Fisher scientific, ND-ONE-W)

96 well plate (VIOLAMO, 1-1601-02)

- 0.5 U/μL Serotropin solution: dissolve Serotropin in 1 mL of 0.6% NaCl solution (supplied).
- 2 U/μL Human Chorionic Gonadotropin (HCG) solution: dissolve HCG in 1.6 mL of 0.6% NaCl solution (supplied).
- 10 mM Tris-EDTA (TE) buffer (pH 7.5): dissolve 8.88 g of Tris-HCl and 3.72 g of EDTA in 1 L of deionized water. Adjust the pH with 5N NaOH.

## 4. Procedure

### 4.1 Preparation of *NEXTi* transgenic plasmid (donor)

*NEXTi* donor plasmids should contain genomic DNA fragments homologous to target sequences (Fig. 1B). While homologous fragments that ranged from 66 bp to 844 bp were used for the donors and resulted in efficient integrations (Table. 1), a longer fragment in the donor resulted in a higher integration rate at least in the case of the *krt12.2.L* locus (Mochii et al., 2024). We usually amplify genomic DNA of 500 – 900 bp including the target 5’UTR and clone it into a vector plasmid using conventional procedures.

1. Clone the DNA fragment including the 5’ UTR sequence of the target gene into a plasmid vector to construct the *NEXTi* donor. For example, an 844 bp fragment containing 698 bp upstream and 146 bp 5′ UTR sequences of *myod1.S* was amplified from the genomic DNA using a high-fidelity polymerase (e.g., PrimeSTAR Max) and cloned into a promoter-less *egfp* plasmid (Suzuki et al., 2010) by standard plasmid construction methods (Mochii et al., 2024) (See Tip 3).

2. Purify the DNA using a standard plasmid purification kit (e.g., PureYield Plasmid System, Promega).

3. (Optional) Extract the DNA with phenol-chloroform and precipitate it with ethanol. Dissolve the DNA in RNase-free TE buffer at 0.5 –1.0 µg/µL (See Tip 4).

1. Dilute the DNA with 1 x Cas 9 buffer at 50 ng /µL, divide it into small aliquots (5 – 10 µL) and store them at –20°C for working stocks.

### 4.2 Synthesis of sgRNA

5. The Cas9 targets should be selected within the 5’ UTR sequence cloned into the donor DNA (See Tip5). A PCR-based strategy was used for sgRNA preparation essentially as previously described (Nakayama et al., 2013; Sakane et al., 2017).

6. Prepare the sequence-specific 5’ oligonucleotides (e.g., for two target sequences of *myod1.S* 5’UTR) and the common 3’ oligonucleotide.

*myod1.S 5*’oligo 1:

5’- TAATACGACTCACTATAGGGGCAAGGTGACTGTGCGTG TTTTAGAGCTAGAAATAGCAAG -3’

*myod1.S 5*’ oligo 2:

5’- TAATACGACTCACTATAGGTTAAAAGATCCACAGCTCG TTTTAGAGCTAGAAATAGCAAG -3’

Common 3’ oligo:

5’- AAAAGCACCGACTCGGTGCCACTTTTTCAAGTTGATAACGGACTAGCCTTATTTTAACTTGCTATTTCTAGCTCTAAAAC -3’

7 Amplify the template DNA by PCR using the oligonucleotides.

#### PCR master mix

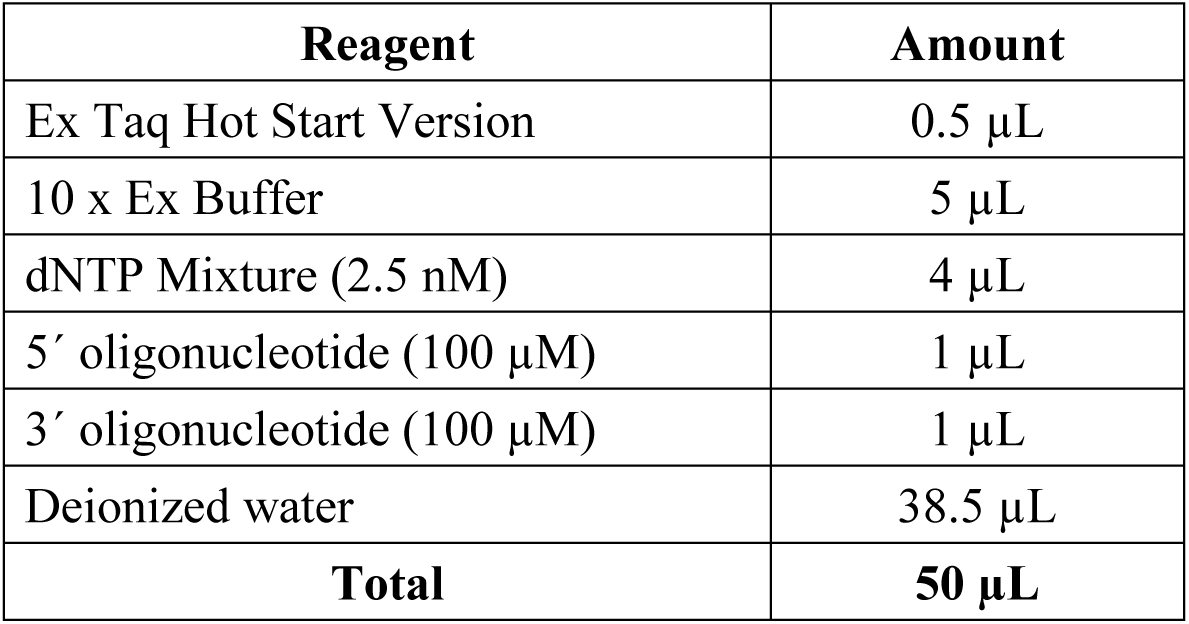

#### PCR cycling conditions

| Step | Temperature | Time | Cycles |
| --- | --- | --- | --- |
| Initial Denaturation | 98°C | 10 sec | 1 |
| Denaturation | 98°C | 10 sec | 12 |
| Annealing | 60°C | 30 sec |  |
| Extension | 72°C | 15 sec |  |
| Final extension | 72°C | 5 min | 1 |
| Hold | 4°C | forever | 1 |

8. Check if the target PCR product is detected as a single band (117 bp) by standard agarose or polyacrylamide gel electrophoresis.

9. Purify the template DNA using a purification kit (e.g., Wizard SV Gel and PCR clean-up system, Promega, A9282). Adjust the concentration to 0.1 µg/µL.

10. Synthesize and purify the sgRNA by using an *in vitro* transcription kit based on T7 RNA polymerase (e.g., Promega, P1320) following the manufacturer’s instructions.

#### Reaction mixture for the RNA synthesis (Promega, P1320)

| Reagent | Amount |
| --- | --- |
| Template DNA (0.2 µg) | 2 µL |
| RiboMAX™ Express T7 2X Buffer | 10 µL |
| Enzyme Mix, T7 Express | 2 µL |
| Nuclease-free water | 6 µL |
| **Total** | **20 µL** |

a. Incubate the reaction mixture at 37°C for 3 – 4 h.
b. Add 1 µL of RNase-free DNase (1 µL/µL) and incubate at 37°C for 15 min.

Add 115 µL of water and 15 µL of 3M sodium acetate to stop the reaction.
Extract with phenol-chloroform.
Add 150 µL of isopropanol and leave at –20°C for at least 15 min.
Centrifuge at 15,000 rpm for 15min.
Remove the supernatant and add 70% EtOH.
Centrifuge at 15,000 rpm for 15 min.
Carefully remove the 70% EtOH.
Dry up the precipitate.
Dissolve sgRNA with nuclease-free water.
Adjust the concentration to 1 – 2 µg/µL and store at –80°C (See Tip6).

11. Prepare master mix for the sgRNA aliquots.

| Reagent | Amount |
| --- | --- |
| sgRNA | 8 $\mu\text{g}$ |
| 10 x Cas9 buffer | 6.2 $\mu\text{l}$ |
| Water | X $\mu\text{L}$ |
| **Total** | **62 $\mu\text{L}$** |

12. Dispense 6.2 µL of the solution into a tube and store at –80℃ (See Tip7).

### 4.3 In vitro fertilization

Standard procedures for *Xenopus* embryo manipulation should follow the protocols of each laboratory or refer to textbooks (Hoppler and Vize, 2012; Sive et al., 2000). Researchers should obtain approval from the Animal Care and Use Committee in their affiliated institutes and perform experiments in accordance with their guideline. In this study, we obtained approval from the committee of University of Hyogo.

1. Inject 50 U of serotropin solution into 2 – 4 females 2 – 3 days before use. The day before use, inject 500 U of HGC solution and keep females at 18°C.
2. Sperm preparation

a. On the day of injection, anesthetize a male in a water tank of 0.2% MS222.
b. After euthanasia, isolate testes and remove the fat body with a pair of fine forceps under a stereomicroscope.
c. Keep the isolated testes in 1 x MMR containing kanamycin on ice (See Tip8).
d. Put a third or half of the testes into a 1.5 mL tube with 1 mL of 0.5 x MMR.
e. Mince the testis fragment in the tube using an ophthalmic scissor.
3. Collect 300–500 eggs by gently squeezing the females and keep them in separate sterile Petri dishes.
4. Add the sperm suspension onto the eggs, mix and spread the eggs gently using a pipet tip.
5. Remove as much of the liquid as possible from the dish and leave the eggs for 5min at 11°C (See Tip9).
6. Soak the dish with 0.1 x MMR for 15 min at 11°C (See Tip10).
7. De-jelly the embryos with de-jellying solution containing L-cysteine for 10 – 15 min (See Tip11).
8. Rinse the embryos with 0.1 x MMR several times.
9. Keep the embryos at a low temperature (11°C) to delay the first cleavage.

### 4.4 Injection

1. Dilute stock Cas9 protein to 1 µg/µL before use. For example, mix 0.5 µL of Cas9 (10 µg/µL) with 4.5 µL of cold 1 x Cas9 buffer.
2. Prepare injection mixture consisting of the following.

| Reagent | Amount |
| --- | --- |
| sgRNA solution | 6.2 µL |
| donor DNA (50 ng/µL) | 1 µL |
| Cas 9 protein (1 µg/µL) | 2 µL |
| **Total** | **9.2 µL** |

After incubating the mixture at room temperature for 10 minutes, place it on ice.

3. Set a microneedle (See **3.2 Setup**) on the microinjector.

4. Fill the microneedle with 1–2 µL of injection mixture (See Tip 12).

5. Transfer 100 –150 fertilized eggs into the injection plate filled with 3-5% Ficoll-0.5 x MMR. Gently shake the injection plate horizontally to direct the fertilized eggs to have their animal side up.

6. At 1 h after the in vitro fertilization, inject 9.2 nL of injection mixture into the animal hemisphere of each fertilized egg. Complete the injection within 1 h (See Tip 13).

7. Transfer the injected eggs in another dish filled with 3% Ficoll–0.5 x MMR and incubate them at 18°C for 2–4 h.

8. Select normal embryos at 4 or 8 cell stages and transfer them into 0.1 x MMR. Unfertilized eggs and severely damaged embryos should be eliminated (see Tip 14).

9. Incubate the embryos at 18°C for 3–4 days until the embryos develop to the late tailbud or tadpole stage (See Tip15–18).

10. Select the embryos with reporter gene expression (See Tip 19). Determine the embryo stages according to Nieuwkoop and Faber (1994).

### 4.5 Germline transmission

#### 3.5.1 In vitro fertilization

1. Raise knock-in frogs to sexual maturation.
2. Outcross matured knock-in animals with wild-type male or female by in vitro fertilization.

### 4.6 Genotyping

1. Deeply anesthetize F1 embryos or tadpoles with MS222 and euthanatize them for genomic DNA extraction.
2. Lyse them in 10 mM Tris-HCl (pH 8.0)-5 mM EDTA-0.5% SDS and add RNase A (10 µg/µL), then incubate at 37°C for 1 h.
3. Treat with Proteinase K (100 µg/µK) at 50°C for 3 h.
4. Extract DNA with phenol and precipitate with ethanol (See Tip 20).
5. Dissolve the DNA in TE buffer.
6. Amplify the junction sequence by using appropriate primer sets (See Tip 21).
7. Purify the PCR product with PCR purification system (e.g., Wizard SV Gel and PCR clean-up system, Promega, A9282) and sequence it.

## 5. Tips

1. The injection mixture is viscous. If the diameter of the needle tip is too small, the needle may become clogged with fluid, and it may not be possible to inject a sufficient volume of the mixture into the eggs. On the other hand, a needle with too-large a tip could seriously damage the injected embryos. We recommend a needle tip with an outer diameter of about 25 µm. The shape of the needle tip is also critical, as a needle with a flat tip damage embryos. The needle tip should be sharpened by breaking it with a pair of fine forceps or hitting against the edge of a glass slide. An Example of the needle tip is shown in Fig.S1.
2. Regarding use of plastalina, refer to a previous protocol (Ogino et al., 2006). Too much clay will cause the solution to overflow. Fill the Petri dish with the clay to less than half of its height. Coating an injection plate with agarose gel also works well.
3. We use the SliCE method (Okegawa and Motohashi, 2015) to clone the PCR product into the vector. Commercial kits (e.g., In-Fusion HD kit, Takara) are also recommended.
4. The donor DNA should be RNase-free.
5. We usually select two or three candidate sequences within a single target 5’ UTR using the CRISPRscan program (https://www.crisprscan.org, Moreno-Mateos et al., 2015) and prepare multiple sgRNAs for the single target gene. The rate of targeted integration is different among the different sgRNAs (Table 2).
6. Purification of sgRNA using a commercial purification kit is also recommended.
7. The solutions and tubes should be pre-chilled on ice to avoid degradation of the sgRNAs.
8. The isolated testes can be stored for several days at 4°C.
9. Excessive liquid causes floating of the eggs at the next step.
10. If the eggs float, quickly sink them using a pipet. Activation of the eggs is confirmed by the contraction of melanin pigments. Do not use a batch of eggs in which the activation is observed in less than 50% of the eggs, because egg quality is critical for successful injection.
11. Insufficient de-jelling causes aggregation of the eggs or interferes with penetration of the microneedle.
12. Drop 1 – 2 µL of Cas9 solution on a parafilm and suck the solution into a microneedle. To prevent the tip of a microneedle from drying out, do not keep the tip in air.
13. Timing of injection is critical. Injection before 1 h or after 2 h post-fertilization results in reduced integration rate. If the rate of normal development is less than 50% after injection, reduce the injection volume, *e.g.*, to 4.6 nL per embryo. Reduced amount of injected sgRNA may reduce integration efficiency (Table 2).
14. It is important to remove unfertilized eggs and abnormally cleaved embryos which were seriously damaged by the injections. The dead or abnormal embryos exert harmful effects on the development of other embryos in the same dish.
15. Density of the embryos should be kept low. We recommend up to 100 embryos in a 100 mm Petri dish. Remove dead and abnormal embryos, and replace the medium every day.
16. Culturing a single embryo in a 96-well plate increases the rate of normal development after injection. It protects healthy embryos from negative influence of dead embryos sharing the culture medium. In the case of the single embryo culture, medium change is unnecessary until the embryo reaches about NF stage 40.
17. 50-90% of the selected embryos develop into normal tadpoles in the standard condition.
18. Ficoll (3%) or methyl-cellulose (0.3%) in the culture medium increase the rate of normally developed embryos.
19. Ubiquitous or ectopic signals are frequently observed at early stages, especially at neural stage, in embryos injected with some donor constructs. Most of the signals become faint or disappeared by a later stage and are considered to be non-specific, probably due to *egfp* transcription from injected donor DNAs that were not integrated into chromosomes.
20. DNA purification using commercially available kit (*e.g.* Promega, A9282) is also recommended.
21. We usually perform the genomic PCR with primers which amplify 0.5 kb to 3 kb fragments. Primer sequences used in Fig.4 are indicated in Table S1.

## 6. Anticipated results

### 6.1 Generation of founders

The tissue-specific eGFP signals were observed in F0 embryos targeted for *myod1.S* (somite), *sox2.L* (central nerve system), and *krt12.2.L* loci (fin; Fig 2). The rate of embryos expressing the reporter gene in a tissue-specific pattern is varied from ∼2% to ∼13% (Table 1) (Mochii et al., 2024). The rate was different between different sgRNAs used for injection even with the same donor DNA. For example, injection of two different sgRNAs, *myod1.S* sg1 and sg2, resulted in 9.3% and 1.8% eGFP-expressing embryos, respectively (Table 2).

**Fig. 2.**
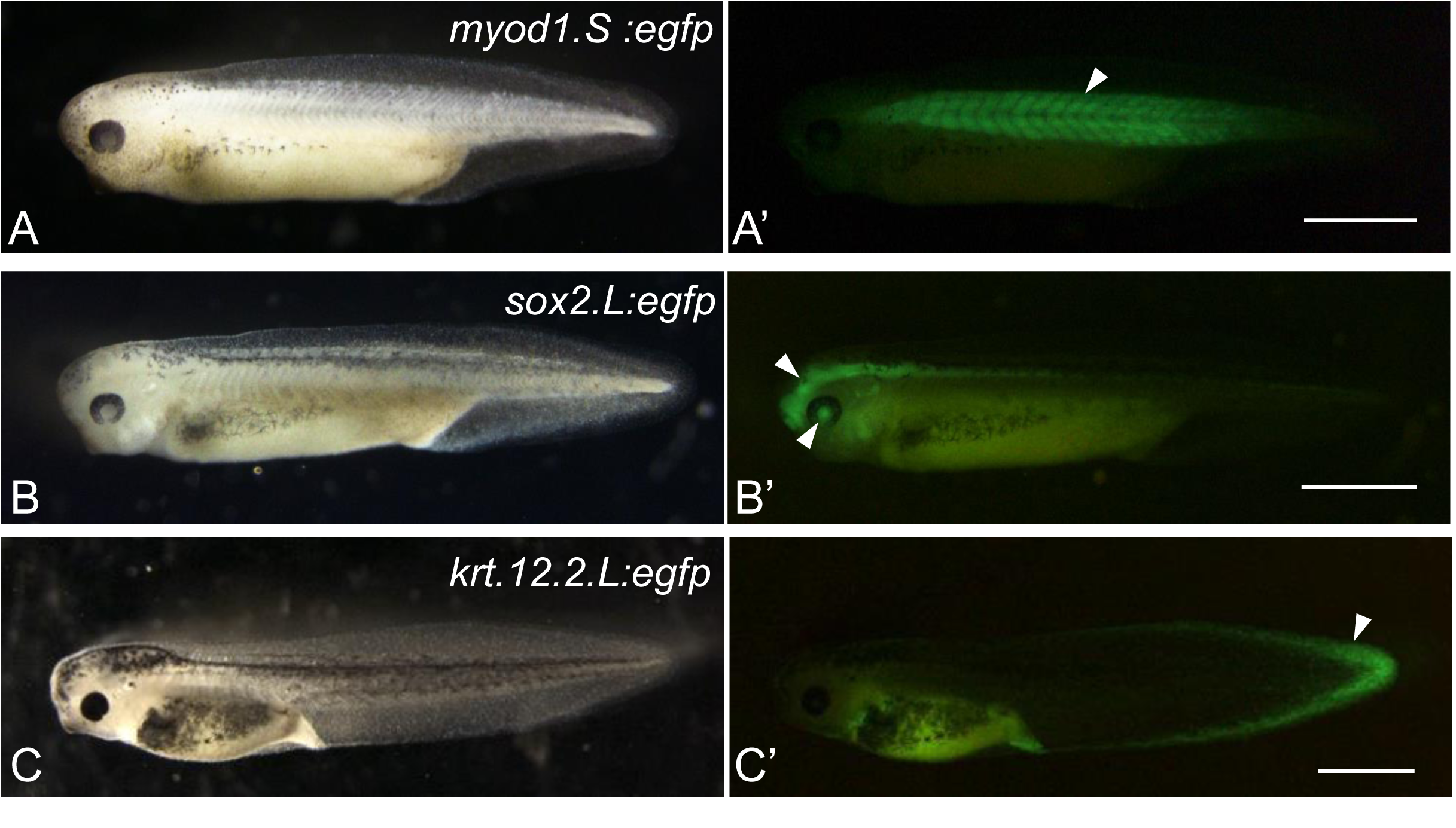
Expression of eGFP in knock-in vector-injected embryos. Representative images of embryos targeted for *myod1.S* (A, A’, NF stage 35/36), *sox2.L* (B, B’, NF stage 37/38) and *krt12.2.L* (C, C’, NF stage 41) loci. (A-C), bright field images. (A’-C’), fluorescence images of (A-C). Clear eGFP signal was observed in somatic muscle (arrowhead in A’), brain and lens (arrowheads in B’), and fin (arrowhead in C’). Scale bar = 1 mm.

**Table 1.** Homologous fragments cloned into donor plasmids and integration rates in founders by *NEXTi*.

| Target locus | Cloned fragment<sup>a</sup><br>(bp) | 5'UTR sequence<sup>b</sup><br>(bp) | Rate of embryos with specific eGFP (%) | Total number of embryos | reference |
| --- | --- | --- | --- | --- | --- |
| *krt12.2.L* | 537 | 66 | 13.2 | 167 | Mochii et.al., 2024 |
|  | 246 | 66 | 6.8 | 161 |  |
|  | 66 | 66 | 3.8 | 261 |  |
| *sox2.L* | 787 | 218 | 2.1 | 1025 | Mochii et.al., 2024 |
| *myod1.S* | 844 | 146 | 6.6 | 708 | Mochii et.al., 2024 |
| *bcan.S* | 489 | 389 | 5.7 | 332 | Mochii et.al., 2024 |

<sup>a</sup>Genomic DNA fragment cloned into donor plasmid. <sup>b</sup> 5'UTR sequence in cloned fragment.

**Table 2.** Integration efficiencies of founders using different sgRNAs or different sgRNA doses.

| Target locus | sgRNA | Amount of sgRNA<br>(pg / embryo) | Rate of embryos with<br>specific eGFP<sup>a</sup> (%) | Total number<br>of embryos | reference |
| --- | --- | --- | --- | --- | --- |
| *myod1.S* | sg1 | 800 | 9.3 | 86 | This work |
|  | sg2 | 800 | 1.8 | 112 |  |
| *myod1.S* | sg1 | 800 | 14.3 | 42 | This work |
|  | sg1 | 400 | 10 | 60 |  |

Amount of donor DNA was constant. <sup>a</sup>Embryos with tissue-specific eGFP expression including mosaic pattern.

### 6.2 Germline transmission of integrated DNA

We confirmed germline transmission of the integrated DNAs by outcrossing matured founders and wild-type frogs by in vitro fertilization. Some F1 embryos of *sox2.L:egfp*, *bcan.S:egfp* and *krt12.2.L:egfp* founders showed eGFP signals in a tissue-specific manner like that observed in their founder embryos (Fig. 3 and data not shown) (Mochii et a., 2024). The rate of F1 embryos showing eGFP signals in a tissue-specific manner varied from 0.3% to 51.6% probably depending on the rate of the knocked-in germ cells in gonads (Table 3). On the other hand, F1 embryos with non-specific eGFP signal were also obtained. In the case of *sox2.L:egfp* founder #2, 2.4% of F1 embryos showed specific eGFP signal in neural tissues (Fig.3A, B), but 3.0% of the siblings showed a weak ubiquitous signal (Fig. 3A, C).

**Fig. 3.**
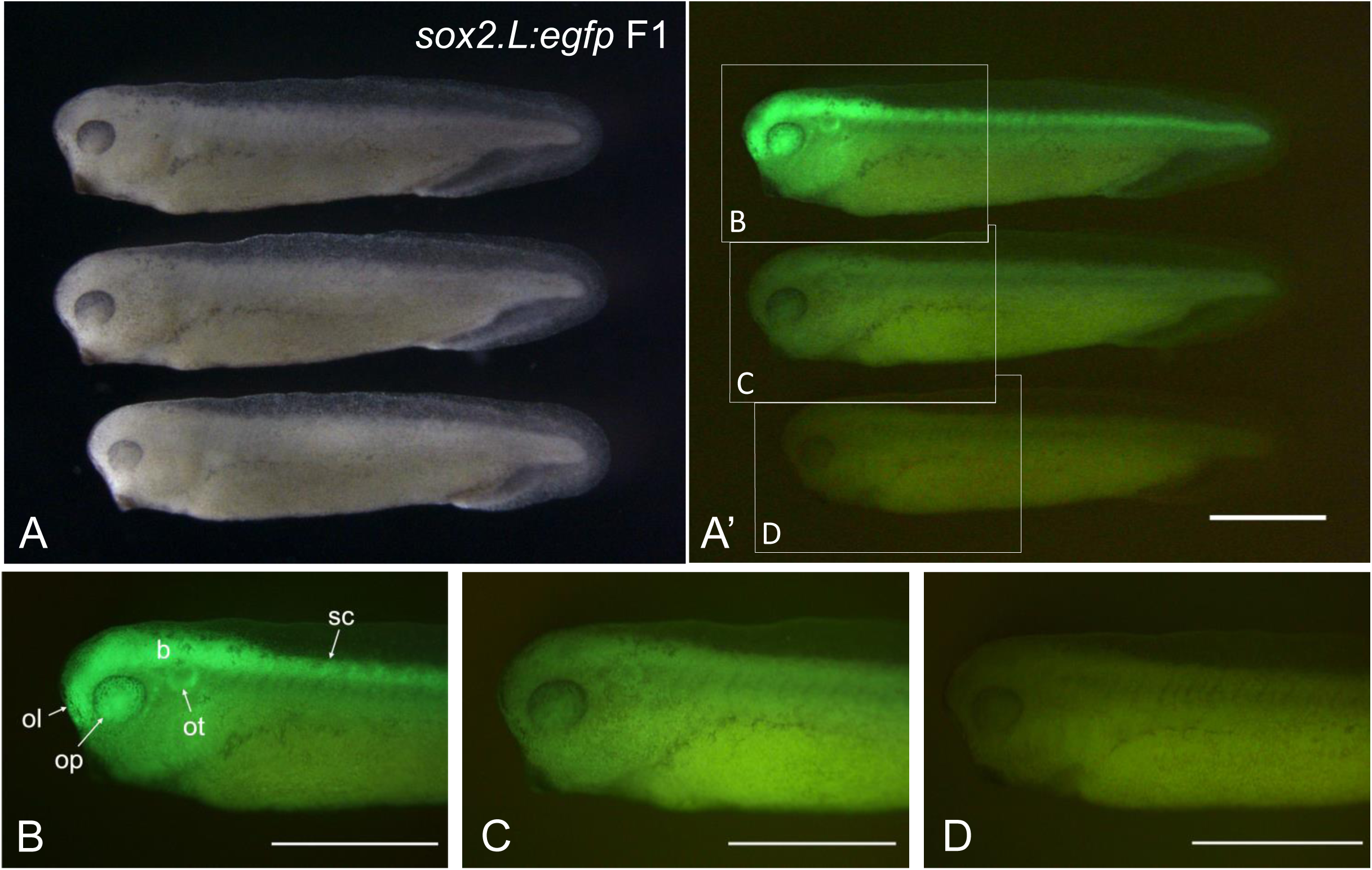
Expression of eGFP in F1 siblings of *sox2.L:egfp*. Representative images of F1 embryos (NF stage 33/34) obtained by crossing a female *sox2.L:egfp* founder #2 (Table. 3) and a wild-type male (A-D). (A) bright field images. (A’), fluorescence image of (A). (B-D) magnified view of each embryo in (A’). (B) specific eGFP signal is observed in brain (b), spinal cord (sc), optic vesicles (op), olfactory pit (ol), and otic vesicles (ot). (C) Ubiquitous eGFP signal at low level is observed. (D) No eGFP signal is observed. Scale bar = 1 mm.

**Fig. 4.**
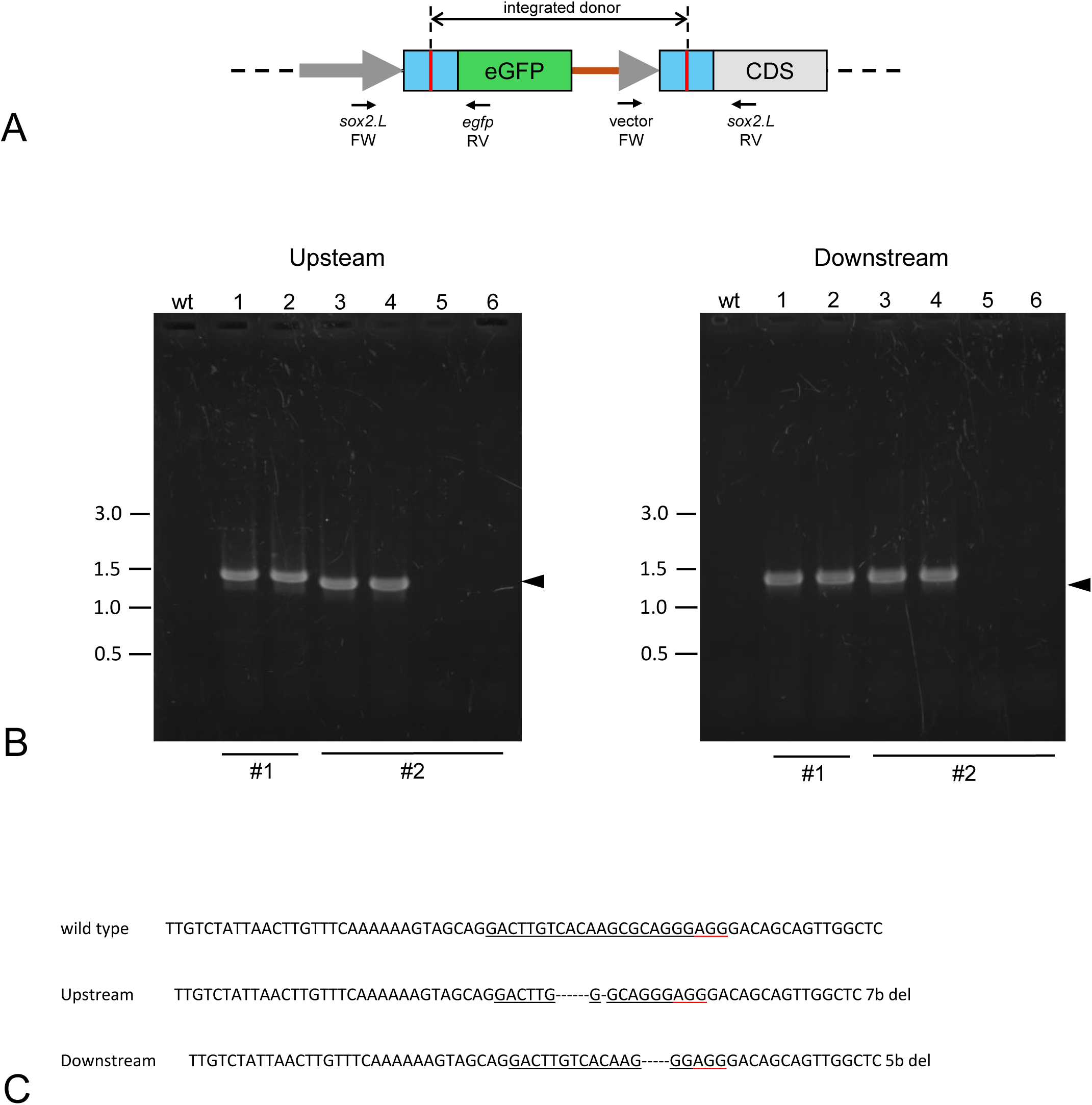
Genotyping of F1 animals. Scheme of an integrated target locus. PCR was performed with genomic DNAs obtained from F1 animals. Upstream and downstream sequences including junctions between the donor and the target regions were amplified using donor vector primers and gene-specific primers. Primer sequences are listed in Table S1. (B) Examples of PCR analysis for *sox2.L:egfp* F1 embryos obtained from the founder #1 (1, 2) and founder #2 (3-6). PCR products for both upstream and downstream junction fragments were observed in DNAs from F1 embryos with specific eGFP expression (1-4) but not from embryos with ubiquitous eGFP (5, 6) or wild-type embryos (wt). Predicted sizes of the upstream and downstream PCR fragments are indicated as arrowheads. PCR products with larger or smaller size indicate insertion or deletion in the junctions. (C) Example of upstream and downstream junction sequences. The PCR products of *sox2.L:egfp* F1 embryo (lane 3) were sequenced directly. Determined sequences are indicated with wild-type partial sequence of the 5’UTR. Underline indicates the sgRNA target sequence with PAM sequence (red underline).

**Table 3.**
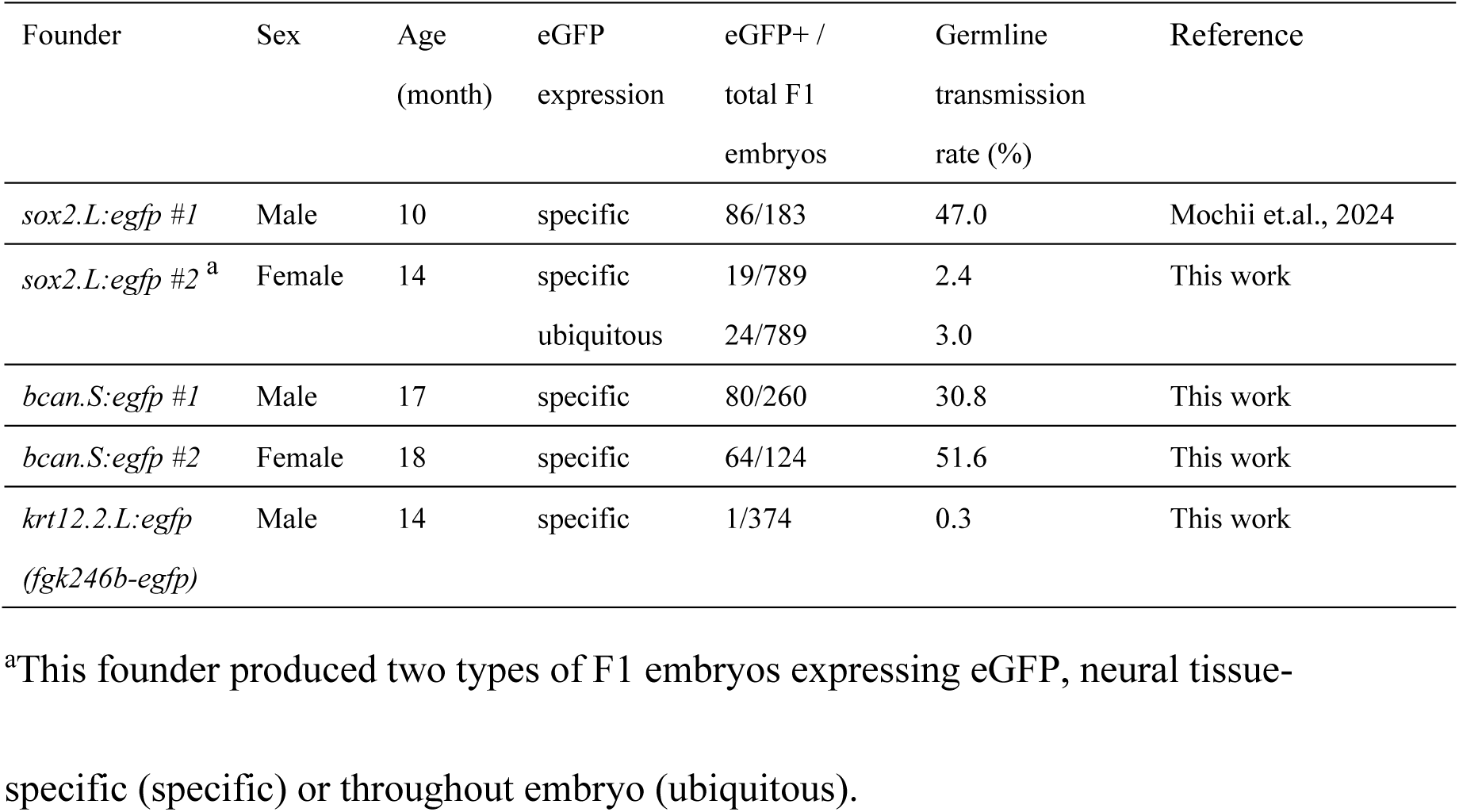
Germline transmission of transgene by *NEXTi*.

Genomic PCR showed the junction between the host genome and donor vector, confirming integration into the 5’UTR of the target loci (Fig. 4). Clear bands were detected in F1 embryos that showed eGFP signal in a tissue-specific manner for both the upstream and downstream junction (Fig. 4B). On the other hand, the PCR band was not detected in either junction in embryos with ubiquitous eGFP signal, suggesting an off-target integration of the donor. The reporter gene integrated in an off-target site may be expressed with low specificity by a possible promoter present in the donor DNA fragment. Sequencing of the amplified fragments showed deletions in both the upstream and downstream junction sites in many cases (Fig. 4C) but insertions in some cases (Mochii et al., 2024).

## Acknowledgments

We thank Dr. Elizabeth Nakajima for critical reading of our manuscript and Dr. Hidefumi Orii for preparation of SLiCE. This research was supported by a JSPS KAKENHI Grant-in-Aid for Scientific Research (C) (JP18K06266 to MM), Grant-in-Aid for Scientific Research (B) (JP21H03829 to KTS), Special grant for research in University of Hyogo (to MM), University of Hyogo Science and Technology Support Foundation (to MM), NIBB Collaborative Research Program (22NIBB331, 23NIBB340 to MM), and JST, CREST, Grant Number JPMJCR 2025 (to KTS), Japan.

---

**Fig. S1.**
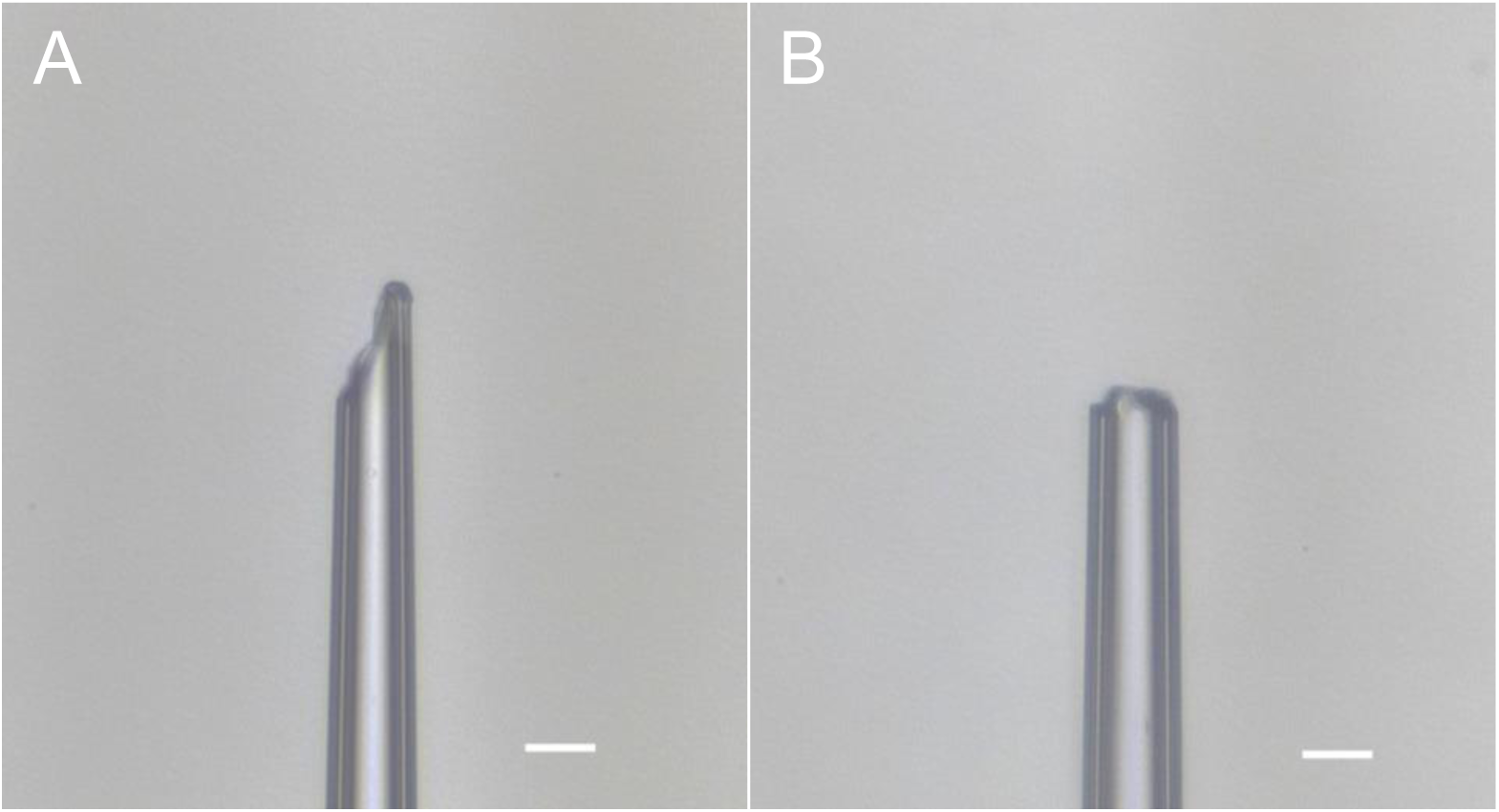
Examples of microneedle tips. The sharp-tipped microneedle (A) is more suitable for the microinjection than the blunt-tipped one (B). Scale bar, 20 µm.

**Table.S1.**
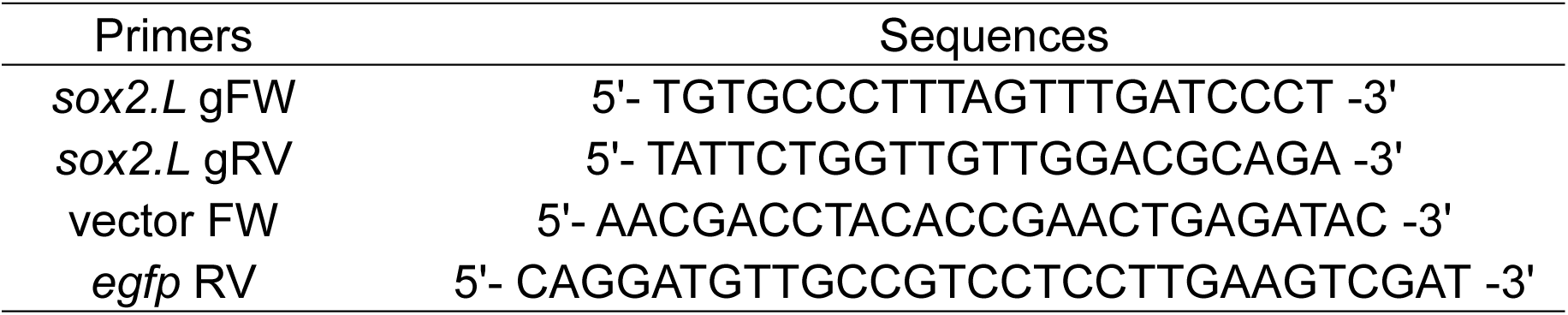
primer sequences for genotyping.

